# Mesodermal FGF and BMP govern the sequential stages of zebrafish thyroid specification

**DOI:** 10.1101/2020.08.13.249540

**Authors:** Benoit Haerlingen, Robert Opitz, Isabelle Vandernoot, Angelo Molinaro, Meghna Shankar, Pierre Gillotay, Achim Trubiroha, Sabine Costagliola

## Abstract

Thyroid tissue is the site for *de novo* synthesis of thyroid hormones which are essential for vertebrate development and growth. Defects in embryonic thyroid morphogenesis are a predominant cause for congenital thyroid diseases but the molecular pathomechanisms are incompletely understood. The first molecularly recognizable step of thyroid development is the specification of thyroid precursors at a defined position in the anterior foregut endoderm. While recent studies identified FGF and BMP pathways as critical signaling factors for thyroid specification, the interplay between extrinsic signaling cues and thyroid transcription factor expression remained elusive. Here, we used zebrafish embryos to decipher the dynamics of thyroid transcription factor induction in relation to FGF and BMP signaling activities in pharyngeal endoderm. We first identified a previously unrecognized endodermal thyroid progenitor cell population expressing Pax2a but not Nkx2.4b. This cell population is characterized by enhanced FGF signaling but initially lacks detectable BMP signaling. A subpopulation of Pax2a-expressing progenitors differentiates subsequently into thyroid lineage-committed precursor cells co-expressing Pax2a and Nkx2.4b. We next combined pharmacological approaches with genetic models permitting inhibition or ectopic overactivation of signaling pathways to timely manipulate FGF and BMP activities. These experiments support a model where FGF signaling primarily regulates Pax2a expression whereas BMP signaling has dual functions in regulation of both Pax2a and Nkx2.4b expression. Collectively, our data allow us to formulate a refined model of thyroid cell specification from foregut endoderm.

## Introduction

The thyroid gland is an essential endocrine tissue regulating key processes in embryonic development and adult homeostasis through the production of thyroid hormones (TH) (Bernal, 2007; Buchholz et al., 2006; Mullur et al., 2014). The functional subunits of thyroid tissue are follicles consisting of a single-layer epithelium of thyroid follicular cells (TFCs), which synthesize THs. Deficiencies in TH synthesis at birth cause congenital hypothyroidism (CH) with about 85% of CH cases being a result of thyroid dysgenesis (TD) but the genetic origin of CH due to TD is known in less than 5% of cases (Wassner, 2018). With respect to the potential etiopathology of TD, it is remarkable that TD cases show an increased prevalence of cardiovascular malformations (Olivieri et al., 2002), implying a possible mechanistic relationship between developmental abnormalities of the heart and the thyroid.

Thyroid morphogenesis has been studied intensively in murine and zebrafish models (reviewed in Nilsson and Fagman, 2017; Opitz et al., 2013; Porazzi et al., 2009). The first molecularly recognizable event of TFC differentiation is the specification of thyroid precursors in an anterior domain of the foregut endoderm. These thyroid lineage-committed precursors are characterized by co-expression of key thyroid transcription factors *Nkx2.1* and *Pax8* in mammals and *nkx2.4b* and *pax2a* in zebrafish (Kimura, 1996; Mansouri et al., 1998; Rohr and Concha, 2000; Wendl et al., 2002). Shortly after specification, the thyroid anlage undergoes a complex series of morphogenetic events including budding, relocalization, folliculogenesis and functional maturation, including the induction of functional differentiation markers such as thyroglobulin (*Tg*), thyroid peroxidase (*Tpo*) or the sodium-iodide symporter (*Slc5a5*) (Alt et al., 2006; Fagman et al., 2006; Fagman and Nilsson, 2011; Opitz et al., 2012; 2011).

Compared to other endoderm-derived organ (Bastidas-Ponce et al., 2017; Ober and Lemaigre, 2018; Zepp and Morrisey, 2019), the molecular mechanisms underlying specification of thyroid precursors are still poorly understood, in particular how extrinsic signaling and intrinsic factors orchestrate the earliest steps of thyroid cell differentiation. Recent studies have highlighted the important roles of fibroblast growth factor (FGF) and bone morphogenetic protein (BMP) signaling. Murine mutant models with defective FGF signaling display severe thyroid hypoplasia or absence of thyroid tissue (Celli et al., 1998; Ohuchi et al., 2000; Revest et al., 2001) and the importance of this pathway for thyroid specification have since been confirmed in various *in vitro* and *in vivo* models (Haerlingen et al., 2019; Kurmann et al., 2015; Longmire et al., 2012; Serls et al., 2005; Wendl et al., 2007). In stem cell-based models, FGF2 stimulation of anterior foregut endoderm cell cultures promotes differentiation of thyroid precursors (Kurmann et al., 2015). Conversely, pharmacological inhibition of FGF signaling blocks thyroid specification in mouse foregut explants and *Xenopus* and zebrafish embryos. In addition, BMP signaling appears to be a crucial factor for early thyroid cell differentiation: BMP4 is a potent stimulator of thyroid precursor differentiation in *in vitro* stem cell models and blockade of BMP signaling during somitogenesis causes a lack of thyroid specification in *Xenopus* and zebrafish embryos (Haerlingen et al., 2019; Kurmann et al., 2015).

Observations made in different species consistently point to precardiac and cardiac mesoderm as a major source of FGF and BMP ligands acting on the foregut endoderm to initiate thyroid cell differentiation (Serls et al., 2005; Vandernoot et al., 2020; Wendl et al., 2007). In all species analyzed, a thyroid anlage is eventually specified in the region of the foregut endoderm that is immediately adjacent to cardiac mesoderm forming the outflow tract structures of the developing heart, and additionally, at the midbrain-hindbrain boundary (MHB) level in zebrafish (Alt et al., 2006; Haerlingen et al., 2019; Norris, 1920; Wendl et al., 2007). Zebrafish studies also showed that loss of cardiac mesoderm or impaired myocardial differentiation is associated with impaired specification of thyroid precursors (Vandernoot et al., 2020) and that perturbed positioning of cardiac mesoderm relative to the prospective thyroid field in the foregut endoderm results in aberrant thyroid development (Haerlingen et al., 2019).

Here, we used zebrafish embryos to spatiotemporally map expression dynamics of key thyroid transcription factors (*pax2a*, *nkx2.4b*) as well as signaling activities of FGF and BMP in the prospective thyroid field of the foregut endoderm. We identified a novel endodermal thyroid progenitor cell population expressing Pax2a and report for the first time that the early thyroid anlage forms by sequential induction of Pax2a and Nkx2.4b. Moreover, we show that this sequence of transcription factor induction mirrors the onset of FGF and BMP signaling activity within the thyroid-forming endoderm. Timed manipulation of FGF and BMP signaling revealed that FGF signaling primarily regulates Pax2a expression whereas BMP signaling has dual functions in regulation of both Pax2a and Nkx2.4b expression. By integrating our data with results from previous studies, we propose an updated model of thyroid cell specification that explains the specific positioning of the thyroid anlage as a result of FGF- and BMP-dependent patterning processes.

## Results

### Sequential induction of thyroid transcription factors Pax2a and Nkx2.4b

According to current models, the differentiation of ventral foregut endodermal cells into thyroid precursors is the earliest recognizable step of thyroid morphogenesis and reportedly occurs at embryonic day 8.5 in mice (co-expression of *Nkx2-1* and *Pax8*) and around 23/24 hpf in zebrafish (co-expression of *nkx2.4b* and *pax2a*) (Fagman et al., 2006; Wendl et al., 2002). While it is commonly accepted that co-expression of these transcription factors is a hallmark of lineage commitment, the induction dynamics and possible molecular mechanisms of thyroid cell specification remained largely unexplored.

To characterize the induction of Pax2a expression in the presumed thyroid field, we performed immunofluorescence (IF) staining (Trubiroha et al., 2018) on a series of wild-type (WT) zebrafish embryos sampled every 2 hours starting at 15/16 hpf. First Pax2a expression in endodermal cells became detectable by 18/19 hpf in cells just ventral to the MHB (Fig. 1A). By 20 hpf, already up to 30 cells with strong Pax2a expression were detectable (Fig. 1B) and their number rapidly increased over the next few hours reaching peak levels at 28 hpf. At this stage, the rostral edge of the Pax2a expression domain was located ventral to the MHB and the domain extended caudally approximately 100 µm along the anterior-posterior axis. Surprisingly, the foregut of 28 hpf embryos contained up to 100 Pax2a+ cells, while the thyroid primordium of 55 hpf zebrafish embryos contains only 30 to 35 differentiated thyroid cells (Trubiroha et al., 2018). Indeed, from 28 hpf onwards, we noticed a progressive decline in the number of Pax2a+ endodermal cells in the thyroid primordium of 55 hpf embryos (see Fig. 1B).

**Fig. 1:**
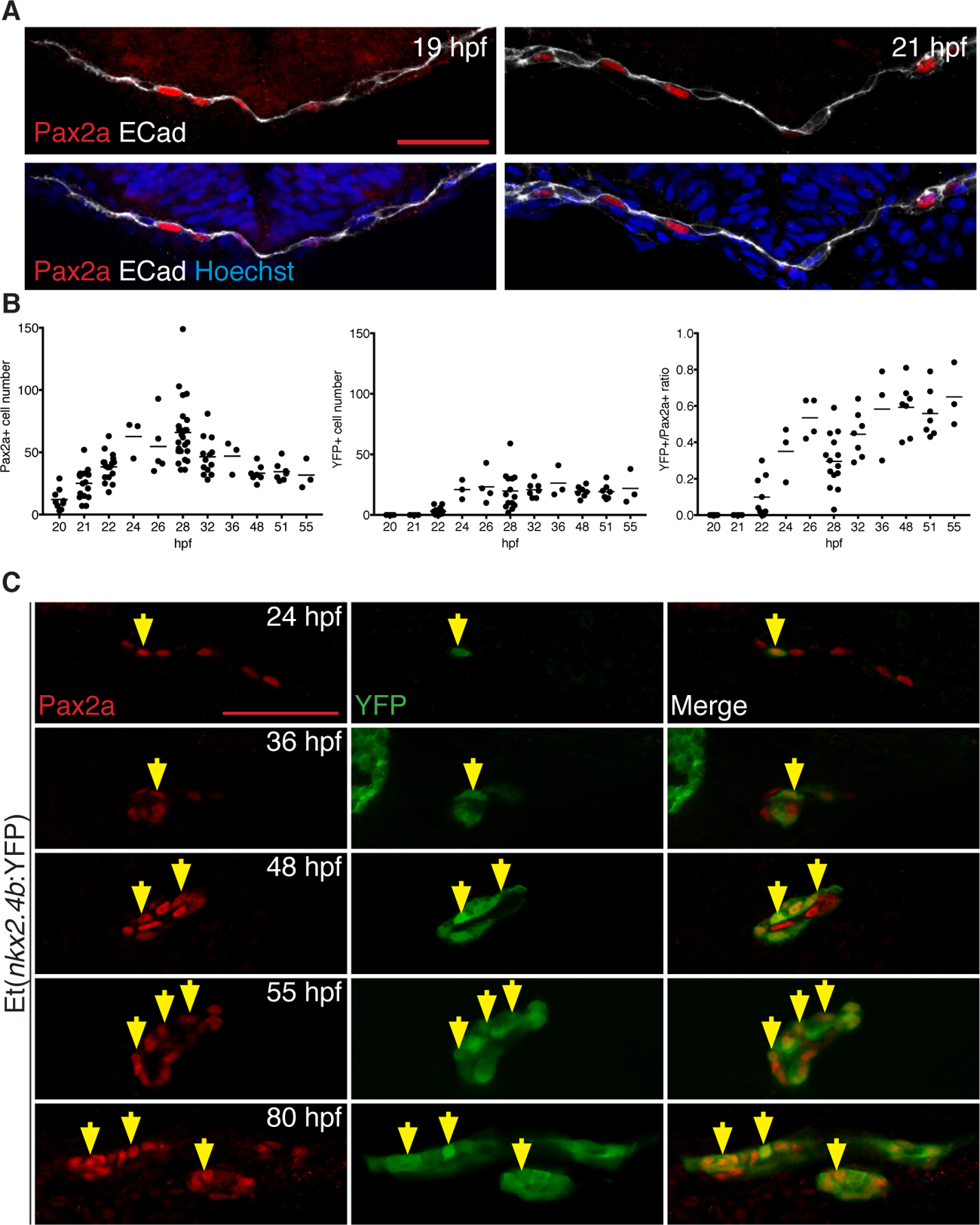
Sequential induction of thyroid transcription factors in foregut endoderm. (A) Pax2a expression in prospective thyroid region of foregut endoderm (labelled by E-Cadherin, Ecad) before thyroid anlage formation. Confocal images of frontal sections. (B) Number of foregut endodermal cells expressing Pax2a (left) and YFP (middle) in Et(*nkx2.4b*:YFP) embryos. Right panel: relative abundance of double positive cells within the Pax2a+ cell population. Dots: value determined in individual embryos; bars: mean values. (C) Pax2a and YFP expression in thyroid region of Et(*nkx2.4b*:YFP) embryos. Confocal images of sagittal (26 to 55 hpf) and coronal sections (80 hpf). Anterior is to the left. Arrows: Pax2a and YFP co-expressing cells. Scale bar: 25 µm.

The concurrent temporal induction of Nkx2.4b expression was monitored using the enhancer trap line Et(*nkx2.4b*:YFP) (Ellingsen et al., 2005), in which the YFP reporter expression neatly correlates with *nkx2.4b* mRNA expression during early zebrafish development (Rohr and Concha, 2000) and [data not shown]. Our IF stainings revealed no detectable YFP expression in the foregut endoderm prior 22 hpf. A faint YFP staining of few endodermal cells was first recognized in some of the 22 hpf embryos analyzed. It was only from 23 hpf onwards that YFP+ cells were reliably detected in the foregut endoderm (Fig. 1B,C) implicating that the onset of *nkx2.4b* expression is delayed by 4-5 hours relative to Pax2a. Quantitative analyses of the YFP+ cell population showed that the number of YFP+ endodermal cells remained stable between 24 and 55 hpf (Fig. 1B).

Co-staining of Pax2a and YFP in Et(*nkx2.4b*:YFP) embryos revealed that *nkx2.4b*:YFP reporter expression is exclusively detectable in cells expressing Pax2a (Fig. 1C), at all analyzed stages. Conversely, only a fraction of Pax2a+ endodermal cells showed co-expression of the *nkx2.4b*:YFP reporter during thyroid anlage formation. The first few endodermal cells co-expressing Pax2a and YFP were detectable at 22 hpf. The ratio of YFP+/Pax2a+ cells fluctuated over time, mainly due to the variations in the number of Pax2a+ cells, but eventually approached a value of one along with the formation of a thyroid primordium composed of thyroid cells co-expressing Pax2a and Nkx2.4b.

In combination, our observations show that zebrafish thyroid specification is in fact a multi-step process that begins several hours earlier than previously reported with the induction of a so far unrecognized population of Pax2a+ endodermal progenitor cells from which only a subpopulation of cells becomes specified into committed Pax2a+/Nkx2.4b+ thyroid precursors in a second step.

### Sequential FGF and BMP signaling activities during thyroid specification

The sequential appearance of Pax2a+ thyroid progenitors and committed Pax2a+/Nkx2.4b+ thyroid precursors raises the question as to possible extrinsic signals regulating these processes. Recently, we showed that FGF and BMP signaling are critical during somitogenesis for regulation of early zebrafish thyroid development (Haerlingen et al., 2019). We therefore used zebrafish biosensor lines to map FGF and BMP pathway activities in the thyroid field.

When using the Tg(*dusp6*:d2EGFP) line (Molina et al., 2007) to monitor cellular FGF signaling activities, first detectable d2EGFP signals in foregut endoderm could be observed in 18/19 hpf embryos (Fig. S1A). We noted that FGF signaling reporter expression in the endoderm was largely confined to Pax2a+ cells and that most Pax2a+ cells showed d2EGFP expression (Fig. S1A). Quantitative analysis showed that about 80% of Pax2a+ thyroid progenitors (n=48) displayed a robust d2EGFP signal (Fig. 2A) linking for the first time the activation of FGF signaling in the foregut endoderm to the expression of the early thyroid marker Pax2a. Pax2a-negative cells with detectable d2EGFP were very rare in foregut endoderm.

**Fig. 2:**
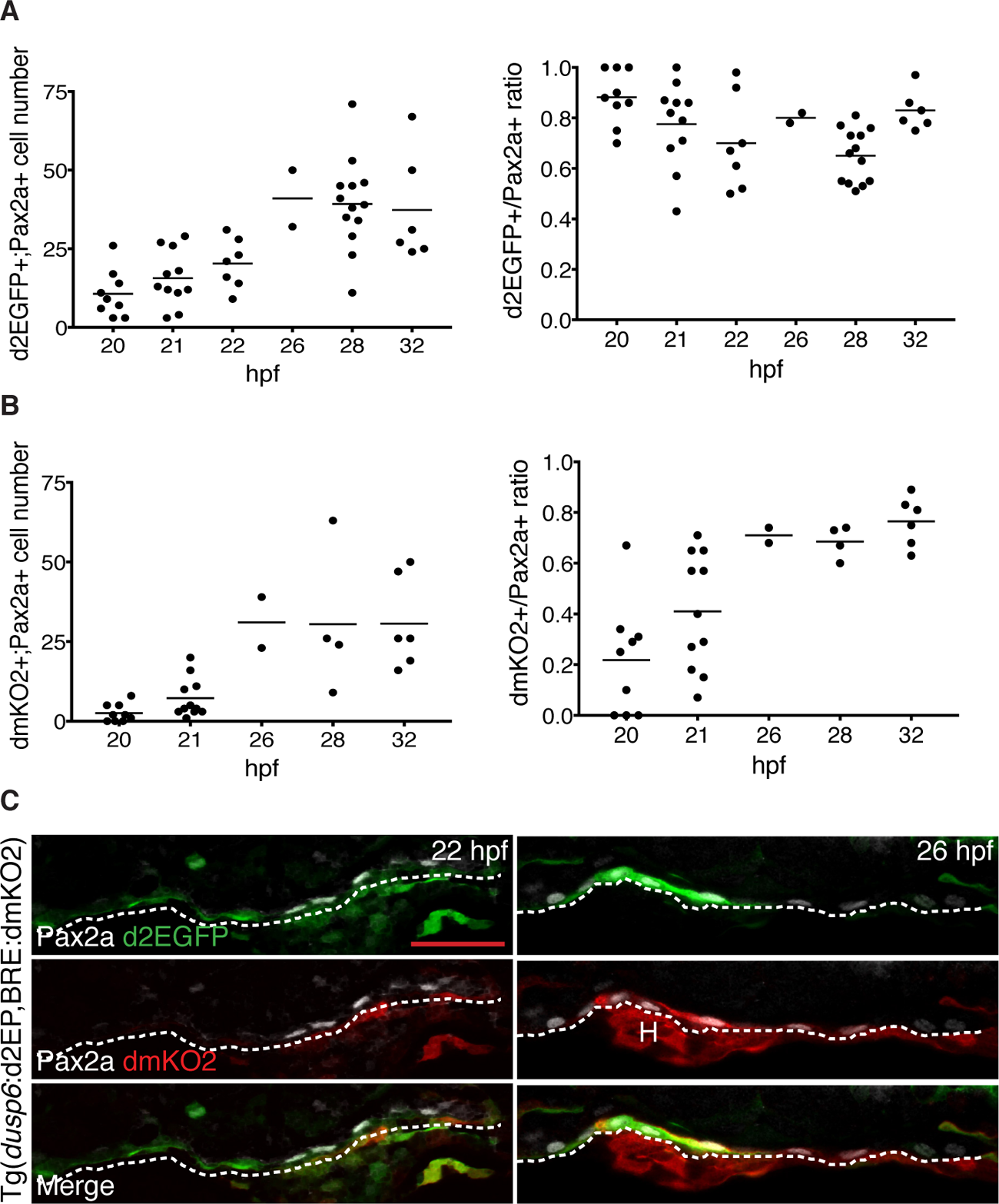
Sequential appearance of FGF and BMP signaling in Pax2-expressing endoderm. (A) FGF signaling reporter expression. Total number of Pax2a+/d2GFP+ double positive cells (left) in foregut endoderm of Tg(dusp6:d2EGFP) embryos and their relative abundance in the Pax2a+ cell population (right). Dots: value determined in individual embryos; bars: mean values. (B) BMP signaling reporter expression. Total number of Pax2a+/dmKO2+ double positive cells (left) in foregut endoderm of Tg(BRE:dmKO2) embryos and their relative abundance in the Pax2a+ cell population (right) (C) Pax2a, d2GFP and dmKO2 expression in thyroid region of Tg(*dusp6*:d2EGFP;BRE:dmKO2) double transgenic embryos. Confocal images of frontal sections. Dashed line: border between endodermal cell layer and ventral foregut mesenchyme. H: heart. Scale bar: 25 µm.

In contrast, when monitoring BMP signaling in the thyroid region using the Tg(BRE:dmKO2) line (Collery and Link, 2011), we found that dmKO2 expression was largely absent in endodermal cells including the Pax2a+ thyroid progenitors before 20 hpf (Fig. S1A). Very few Pax2a+ cells began to express low levels of dmKO2 reporter between 20 and 22 hpf (Fig. 2B). Thus, from 18 to 22 hpf, endodermal cells of the prospective thyroid region showed detectable FGF signaling limited to Pax2a+ thyroid progenitors but lacked detectable BMP signaling activity. This pattern changed from 22 hpf onwards, as we detected an increased number of endodermal cells with dmKO2 reporter expression (Fig. 2B). We noted that, in the foregut endoderm, BMP signaling reporter expression was largely restricted to Pax2a+ cells (Fig. S1A). More importantly, BMP signaling in Pax2a+ cells was linked with up-regulation of Nkx2.4b expression in Pax2a+ cells. Specifically, in double transgenic Tg(BRE:dmKO2),Et(*nkx2.4b*:YFP) embryos, almost all dmKO2+ endodermal cells showed YFP expression and *vice versa* (Fig. S1B). In addition, mapping of FGF and BMP signaling showed the most intense reporter signals in endodermal cells located in close vicinity to cardiac outflow tract (Fig. 2C) supporting the notion of cardiac mesoderm as an important source of FGF and BMP signals for thyroid specification (Vandernoot et al., 2020).In combination, our mapping studies of the thyroid endoderm showed that FGF signaling is a defining characteristic of the early Pax2a+ thyroid progenitor cell population and indicate that the emergence of committed Pax2a+/Nkx2.4b+ thyroid precursors is associated with concurrent induction of BMP signaling in Pax2a+ cells.

### BMP and FGF signaling regulate the thyroid progenitor pool size

Our observations suggested that specific epistatic relationships between FGF signaling and Pax2a expression and between BMP signaling and Nkx2.4b expression might be at the core of the sequential induction of Pax2a+/Nkx2.4b+ thyroid precursors. To address this, we used a small molecule inhibitor approach with a refined temporal treatment schedule to assess how periods sensitive to FGF and BMP inhibition relate to the temporal dynamics of endodermal FGF and BMP signaling.

For this purpose, we treated embryos for short intervals during somitogenesis with DMH1 (12 µM), a small molecule inhibitor of BMP signaling (Haerlingen et al., 2019; Hao et al., 2014). Recapitulating previous results, continuous DMH1 treatment during somitogenesis prevented thyroid specification (Fig. 3 and Fig. S2A). Analysis of thyroid marker expression following shorter DMH1 treatment periods showed that inhibition of BMP signaling from 10-13 hpf was as effective as the long-term DMH1 treatment in blocking early thyroid development. Inhibition of BMP signaling during mid-somitogenesis, i.e. from 13-16 hpf or 16-19 hpf caused only mild effects on thyroid marker expression. However, applying DMH1 just prior to the concurrent onset of BMP signaling reporter and *nkx2.4b* expression, between 19 and 21 hpf, reduced or abolished *nkx2.4b* expression in almost all embryos. Furthermore, blockage of BMP signaling between either 19-21 or 21-24 hpf greatly diminished *tg* expression indicating that BMP signaling during the specification period is critical for normal thyroid development (Fig. 3). Similar short-term treatment experiments with a potent inhibitor of FGF signaling (8 µM PD166866) did not highlight a specific FGF-dependent window for *nkx2.4b* and *tg,* although continuous PD166866 treatment during somitogenesis caused a complete absence of thyroid specification (Fig. 3).

**Fig. 3:**
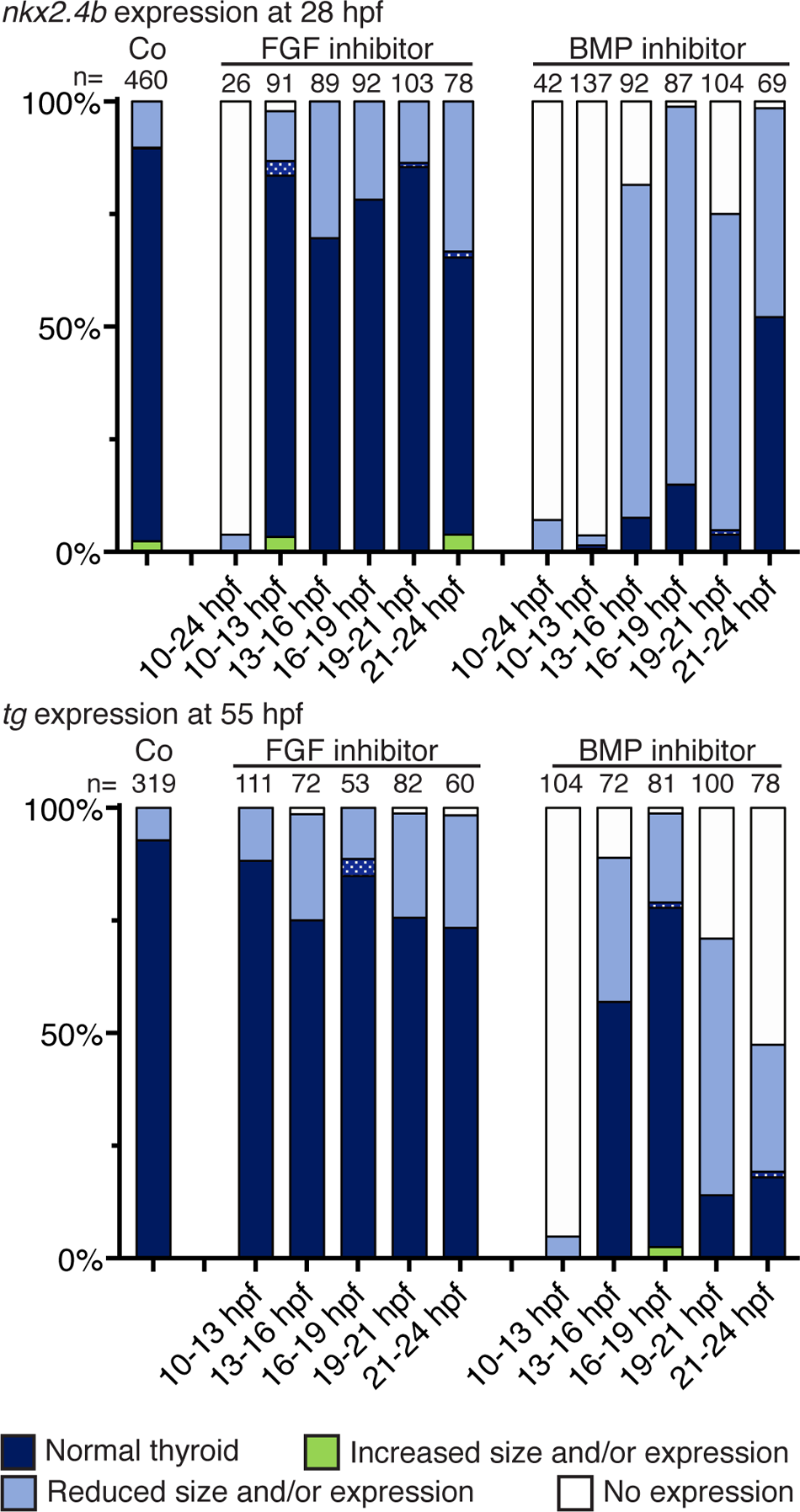
Effects of stage-dependent inhibition of FGF and BMP signaling on thyroid specification Distribution of thyroid phenotypes induced by inhibition of FGF (8 µM PD166866) and BMP signaling (12 µM DMH1). Treatment periods are indicated below each bar. Drugged embryos were analyzed by WISH for *nkx2.4b* (28 hpf) and *tg* expression (55 hpf). Thyroid phenotypes were visually classified into four categories. Dotted pattern: abnormal morphology of thyroid marker expression domain. Co: Control condition (0.1% DMSO).

In addition, global inhibition of BMP signaling during early somitogenesis reduced dramatically the number of Pax2a+ thyroid progenitors at 22 hpf (Fig. S2C) and 28 hpf (Fig. S2B). Suppressive effects of BMP inhibition diminished gradually the later the BMP inhibitor treatment was applied but BMP inhibition just prior to thyroid specification still had a mild to moderate suppressive effect on the total number of Pax2a+ cells. Interestingly, an inverse temporal relationship was observed for FGF signaling (Fig. S2B). Short-term treatment with PD166866 had the strongest suppressive effect on Pax2a expression at late somitogenesis stages whereas transient inhibition of FGF signaling at early somitogenesis did not affect the number of Pax2a+ cells (Fig. S2C).

Integrating these data with results from our mapping experiments, we conclude that periods where Pax2a expression was sensitive to FGF inhibitor treatments correlated closely with the temporal evolution of FGF signaling in foregut endodermal cells. Such a correlation was less evident for BMP signaling as global BMP inhibition evoked the strongest effects on Pax2a expression during periods when BMP signaling specifically in the foregut endoderm was barely detectable. Since BMP signaling plays critical roles in mesoderm patterning and differentiation (Prummel et al., 2020; Row et al., 2018), we therefore hypothesize that the suppressive effects of early global BMP inhibition might stem from impaired patterning of the foregut mesenchyme (including the cardiogenic LPM) rather than from direct effects on endoderm.

To address the latter point, we treated embryos of the Tg(*dusp6*:d2EGFP) reporter line with DMH1 and analyzed d2EGFP reporter expression in foregut endoderm of 22 hpf embryos (Fig. S3). Interestingly, inhibition of BMP signaling during early and mid-somitogenesis stages not only reduced the number of Pax2a+ endodermal cells showing d2EGFP reporter expression but also greatly reduced overall FGF signaling reporter expression in the endoderm, compared to control embryos (Fig. S3B). In contrast, mesenchymal FGF signaling reporter expression was still detectable in DMH1-treated embryos.

### Overactivation of FGF signaling enlarges the pool of Pax2a+ thyroid progenitors

We next investigated if and when enhanced FGF signaling alter thyroid cell specification by using a transgenic zebrafish line Tg(*hsp70*:ca-fgfr1) with a heat shock-inducible expression cassette for a constitutively active Fgfr1 (Marques et al., 2008). Heat shock (HS) induction of FGF signaling at 10 hpf barely affected thyroid specification (Fig. 4A). However, overactivation of FGF signaling at mid (15 hpf) and late somitogenesis stages (20 hpf) resulted in enhanced *nkx2.4b* expression and occasionally in a caudal expansion of the *nkx2.4b* expression domain in 28 hpf embryos (Fig. 4A,B). Genotyping of a portion of stained embryos confirmed that all embryos presenting with enhanced *nkx2.4b* staining carried the HS-inducible transgene.

**Fig. 4:**
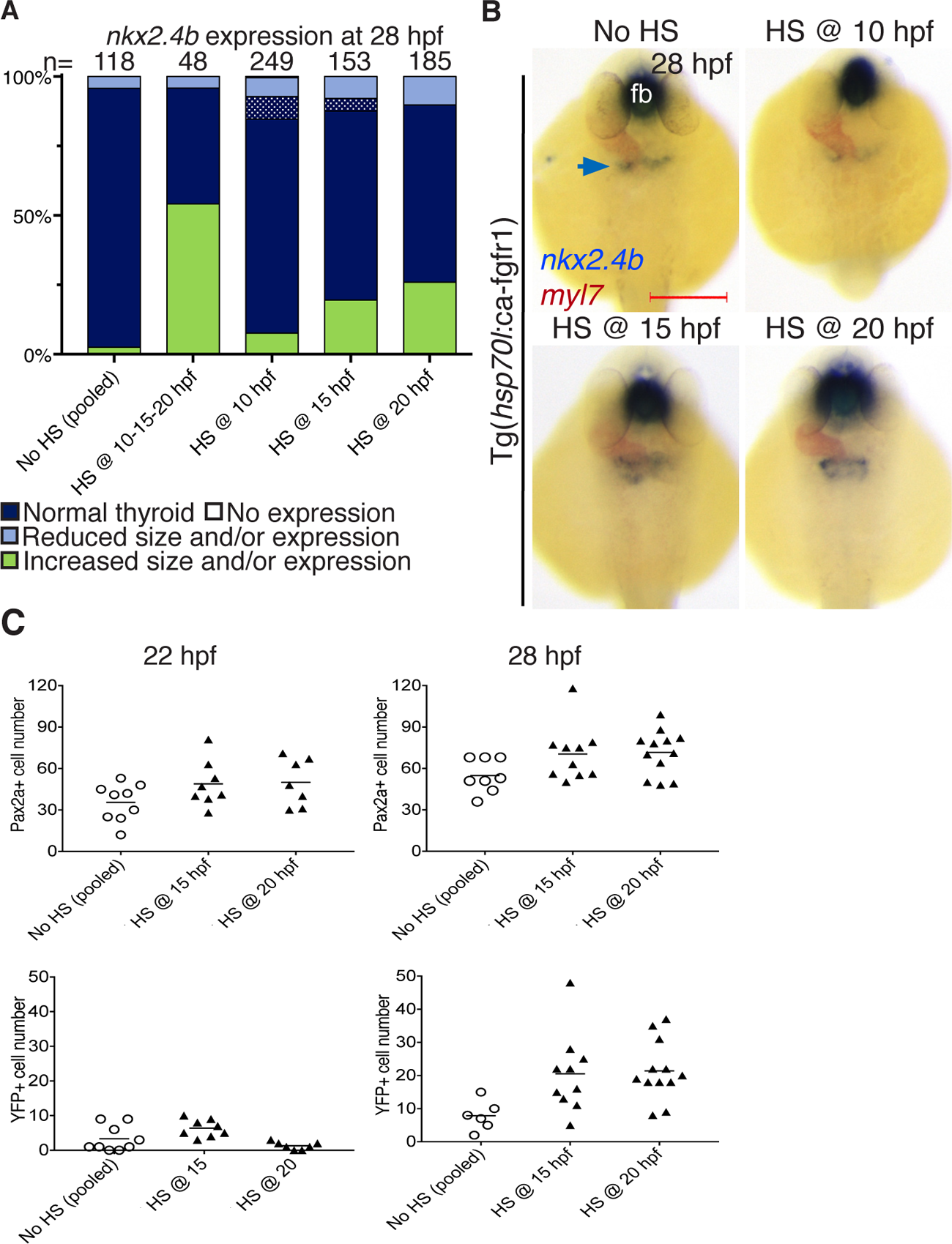
Enhanced FGF signaling during late somitogenesis promotes thyroid precursor specification. (A) Distribution of thyroid phenotypes (*nkx2.4b* expression at 28 hpf) induced by overactivation of FGF signaling by HS of Tg(*hsp70l*:ca-fgfr1) embryos. Phenotypic data shown include carriers and non-carriers of the HS-inducible transgene. Control condition consists of pooled data from non-heat shocked siblings from each of the HS experiments. Dotted pattern: abnormal morphology of thyroid marker expression domain. (B) Major thyroid phenotypes recovered at 28 hpf by *in situ* hybridization of *nkx2.4b* and *myl7* (cardiomyocytes) following HS-induced over-activation of FGF signaling. Dorsal views, anterior to the top. fb: forebrain. Scale bar: 200 µm. (C) Total number of foregut endoderm cells expressing Pax2a (upper panels) and YFP (lower panels) following HS treatment of Tg(*hsp70l*:ca-fgfr1);Et(*nkx2.4b*:YFP) double transgenic embryos at indicated time points. Cell numbers were determined for 22 and 28 hpf embryos. Dots: value determined in individual embryos; bars: mean values. Data include carriers and non-carriers of the HS-inducible transgene.

We also analyzed the effects of such overactivations of FGF signaling on the Pax2a+ thyroid progenitors and observed an increase in the number of Pax2a+ endodermal cells in 22 and 28 hpf embryos (Fig. 4C). Although technical limitations (stained tissue mounted firmly on glass slides) prevented us from using PCR-based genotyping to delineate the genotype of individual embryos, we observed robust shifts towards higher Pax2a+ cell number following HS-induced overactivation of FGF signaling.

Importantly, overactivation of FGF signaling in double transgenic Tg(*hsp70*:ca-fgfr1);Et(*nkx2.4b*:YFP) embryos caused an increase in the number of YFP+ thyroid precursors at 28 hpf (Fig. 4C) but did not result in their premature differentiation(Fig. 4C). In combination, these results suggest that FGF signaling primarily regulates the size of the Pax2a+ progenitor population thereby indirectly enhancing the number of Nkx2.4b+ thyroid precursors differentiating from an enlarged progenitor pool.

### Overactivation of BMP signaling induces supernumerary thyroid precursors

Our observations that Nkx2.4b induction in Pax2a+ progenitors occurs concurrent with up-regulated BMP signaling in a subpopulation of Pax2a+ cells suggests that Nkx2.4b induction is BMP-dependent. To address this, we examined if enhanced BMP signaling prior to and/or during the period of *nkx2.4b* induction affects the specification of Pax2a+/Nkx2.4+ thyroid precursors. For that, we crossed the Nkx2.4b reporter line Et(*nkx2.4b*:YFP) with the transgenic line Tg(*hsp70*:*bmp2b*) carrying a HS-inducible expression cassette for zebrafish Bmp2b (Chocron et al., 2007).

HS treatments performed shortly before (15 hpf) or at the initiation of thyroid specification (20 hpf) increased the size and enhanced the staining intensity of the *nkx2.4b* expression domain of 28 hpf embryos (Fig. 5A,B). Genotyping confirmed that embryos showing enhanced *nkx2.4b* staining carried the HS-inducible transgene. Overactivation of BMP signaling during mid and late somitogenesis did not cause overt effects on foregut morphogenesis, as judged from WISH staining of foregut markers (Fig. 5B) and analyses of GFP reporter expression in Tg(*sox17*:EGFP) embryos (data not shown).

**Fig. 5:**
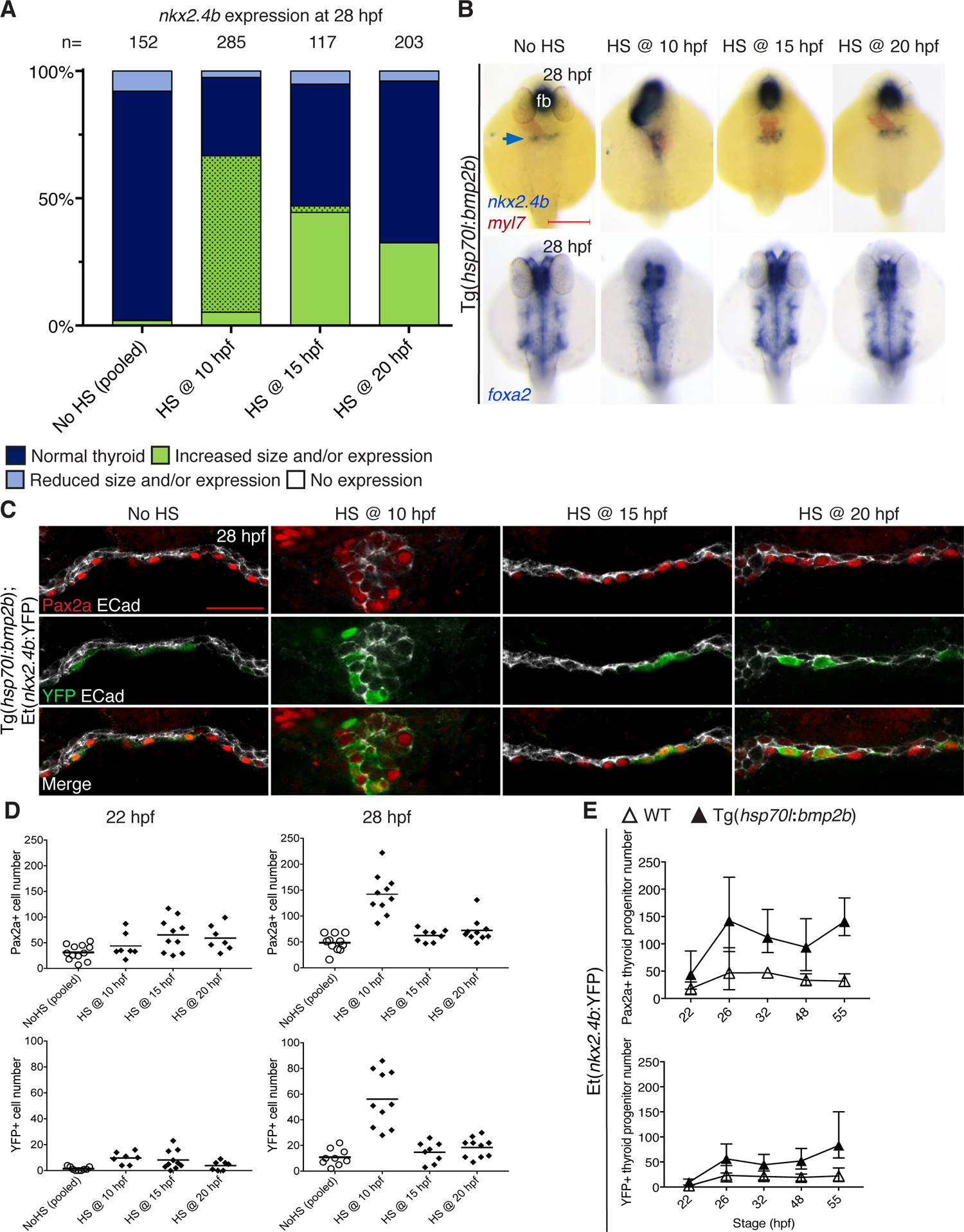
Enhanced BMP signaling promotes thyroid precursor specification. (A) Distribution of thyroid phenotypes (*nkx2.4b* expression at 28 hpf) recovered after HS-induced BMP signaling overactivation in Tg(*hsp70l*:*bmp2b*) embryos. Phenotypic data include carriers and non-carriers of the HS-inducible transgene. Control condition consists of pooled data from non-heat shocked siblings from each of the HS experiments. Dotted pattern: abnormal morphology of thyroid marker expression domains. (B) Thyroid (upper panels) and foregut (lower panels) phenotypes recovered at 28 hpf by *in situ* hybridization following HS-induced over-activation of BMP signaling. Dorsal views, anterior to the top. fb: forebrain. Scale bar: 200 µm. (C) Pax2a, YFP and E-cadherin (ECad) expression in thyroid region of Tg(*hsp70l*:*bmp2b*);Et(*nkx2.4b*:YFP) double transgenic embryos at 28 hpf. Confocal images of frontal sections. Note the severely perturbed morphology of the foregut endoderm following HS treatment at 10 hpf. Scale bar: 25 µm. (D) Total number of foregut endoderm cells expressing Pax2a (upper panels) and YFP (lower panels) following HS treatment of Tg(*hsp70l*:*bmp2b*);Et(*nkx2.4b*:YFP) double transgenic embryos at indicated time points. Cell numbers were determined for 22 and 28 hpf embryos. Dots: value determined in individual embryos; bars: mean values. Data include carriers and non-carriers of the HS-inducible transgene. (E) Number of foregut endoderm cells expressing Pax2a (upper panel) and YFP (lower panel) in embryos at the indicated stages following a single HS treatment of Tg(*hsp70l*:*bmp2b*);Et(*nkx2.4b*:YFP) embryos at 10 hpf. Mean values and standard deviations are shown.

Additional confocal analyses of IF-stained embryos (heat-shocked at either 15 or 20 hpf) revealed an increased number of Pax2a+/Nkx2.4b+ thyroid precursors at both 22 and 28 hpf in a morphologically intact foregut endoderm (Fig. 5D). While these findings are in line with the notion that BMP signaling promotes *nkx2.4b* induction, we also observed that enhanced BMP signaling caused a more rapid induction of Pax2a expression in foregut endoderm (Fig. 5D). Total numbers of Pax2a+ cells were increased at 22 hpf whereas such a surplus of Pax2a+ cells was less evident at later stages (Fig. 5D) when Pax2a+ cell numbers are peaking in normally developing embryos (see Fig. 1B). It thus appears that enhanced BMP signaling at mid and late somitogenesis stages accelerated the onset of Pax2a+ cells but had modest effects on the final pool size of thyroid progenitors. Another key observation from these HS experiments was that enhanced BMP signaling failed to induce Nkx2.4b in all Pax2a+ cells so that Nkx2.4b expression was still restricted to only a fraction of endodermal Pax2a+ cells. Collectively, these experiments showed that ectopic overactivation of BMP signaling prior to and during the onset of thyroid specification affected both Pax2a and Nkx2.4b expression dynamics promoting the differentiation of supernumerary thyroid precursors.

Our BMP overactivation experiments also addressed thyroid specification in embryos heat-shocked at early somitogenesis (HS at 10 hpf) as short-term inhibition of BMP signaling at this stage caused a complete loss of thyroid specification (see Fig. 3).When analyzing thyroid development in Tg(*hsp70*:*bmp2b*) embryos heat-shocked at 10 hpf, we observed the most severe thyroid phenotypes of our HS series. In 28 hpf embryos from this specific experimental group, the thyroid anlage was greatly enlarged and showed a marked caudal expansion along the midline, flanked bilaterally by elongated strands of *myl7*-expressing cardiac mesoderm (Fig. 5B and Fig. S4). Expression of the foregut endoderm marker *foxa2* (Fig. 5B) and GFP reporter in Tg(*sox17*:EGFP) embryos (Fig. S4B) revealed a severely perturbed morphology of the prospective pharyngeal endoderm due to overactivation of BMP signaling. Confocal imaging showed that instead of a sheet-like morphology, the foregut endoderm had a rod-like morphology along its entire extension in embryos heat-shocked at 10 hpf (Fig. 5C and Fig. S4C). Both Pax2a+/Nkx2.4b-progenitors and Pax2a+/Nkx2.4b+ thyroid precursors were intermingled along the dorsoventral axis and over far stretches along the AP axis of these rod-like endodermal structures (Fig. 5C).

Quantitative analyses of 28 hpf embryos confirmed an at least 3-fold increase in the total number of Pax2a-expressing cells and an up to 5-fold increase of Pax2a+/Nkx2.4b+ thyroid precursors following HS of Tg(*hsp70*:*bmp2b*) embryos at 10 hpf (Fig. 5D). Interestingly, while Pax2a+/Nkx2.4b+ thyroid precursors are normally very low in number at 22 hpf, BMP overactivation increased their number more than 6-fold (Fig. 5D). Yet, the absence of detectable YFP expression at 19 hpf suggests that there was no precocious but a more rapid induction of thyroid markers under these conditions (data not shown). Conversely, we observed that the rapid onset of thyroid precursor specification resulted in a permanent amplification of the pool of thyroid precursors so that the size of the thyroid cell population at 55 hpf is on average at least 3 times higher compared to control embryos (Fig. 5E and Fig. S4C). Thus, in addition to the period of thyroid specification at late somitogenesis, our experimental manipulations of BMP signaling identified early somitogenesis as another period where BMP signaling is critical for thyroid specification.

### Combinatorial FGF and BMP signaling is required for thyroid specification

Our pathway manipulations attributed specific and overlapping roles of FGF and BMP signaling on the induction of Pax2a and Nkx2.4b. To assess the effects of combinatorial overactivation of FGF and BMP signalings, we generated Tg(*hsp70*:ca-fgfr1;*hsp70*:*bmp2b*) double transgenic embryos and performed HS experiments at 10, 15 or 20 hpf. Thyroid phenotypes recovered in double transgenic embryos heat-shocked at 10 hpf by WISH staining of *nkx2.4b* at 28 hpf were similar to those observed following timed overactivation of BMP signaling (see Fig. 5B).). However, for HS performed at 15 and 20 hpf, we observed that combined FGF and BMP overactivation resulted in a caudal expansion of the *nkx2.4b* expression domain with enhanced expression, compared to overactivation of a single signaling pathway (Fig. 6 and Fig. S5). This suggests that combined FGF and BMP activation is required during late somitogenesis to promote full thyroid specification.

**Fig. 6:**
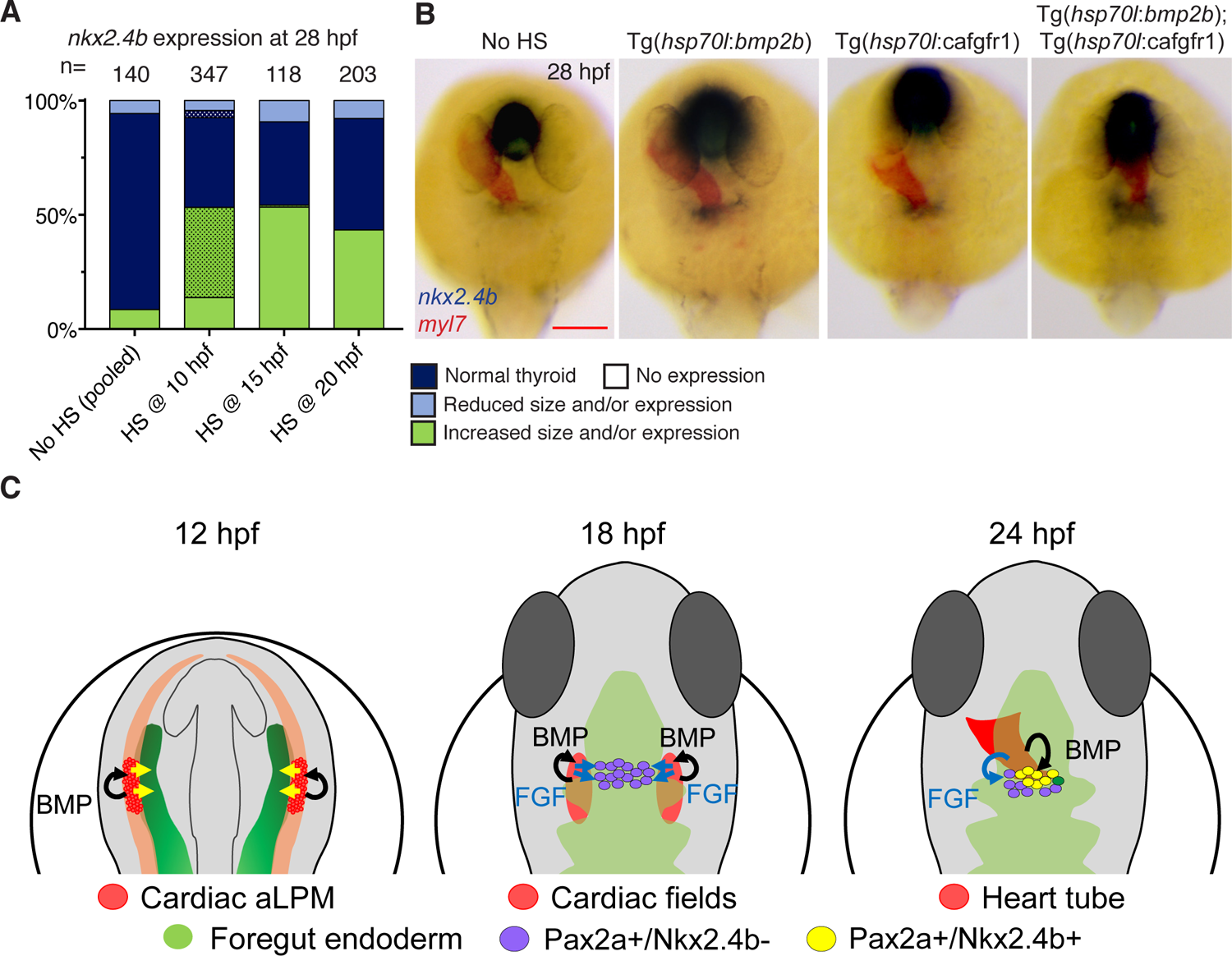
Cooperative effects of FGF and BMP on thyroid specification. (A) Distribution of thyroid phenotypes (*nkx2.4b* expression at 28 hpf) recovered following combined HS-induced BMP and FGF signaling overactivation in Tg(*hsp70l*:*bmp2b; hsp70l*:ca-fgfr1) embryos. Data shown include carriers and non-carriers of the HS-inducible transgenes. Genotyping of a subset of embryos confirmed that promoting effects of 10 hpf HS treatment on *nkx2.4b* expression are exclusively present in embryos carrying the Tg(*hsp70l*:*bmp2b*) transgene. For HS treatments performed at 15 and 20 hpf, activation of either FGF or BMP signaling had promoting effects on *nkx2.4b* expression. Control condition consists of pooled data from non-heat shocked embryos collected during each of the HS experiments. Dotted pattern: abnormal morphology of thyroid marker expression domains. (B) Major thyroid phenotypes recovered at 28 hpf by *in situ* hybridization following HS-treatment at 20 hpf. Dorsal views, anterior to the top. fb: forebrain. Scale bar: 200 µm. (C) Refined model of thyroid precursor specification. Early during somitogenesis, the anterior foregut endoderm (green) receives patterning signals (yellow arrows) from anterior lateral plate mesoderm (aLPM). At around 18 hpf, FGF signals (blue arrows) derived from adjacent cardiac mesoderm (red) induce Pax2a expression in endodermal cells. Up to this stage, active BMP signaling is limited to mesodermal cells. Around 23/24 hpf, a subpopulation of Pax2a+ endodermal cells, located closest to the cardiac outflow tract mesoderm, receives BMP signals inducing Nkx2.4b expression. Pax2a+/Nkx2.4b+ cells represent committed thyroid cell precursors while Pax2a+/Nkx2.4b-cells assume alternative cell fates.

We next examined if activity of both pathways is strictly required for thyroid specification or if overactivation of one pathway can compensate for the loss of the other. In a first experiment, we repeatedly heat-shocked Tg(*hsp70l*:ca-fgfr1) and Tg(*hsp70l*:*bmp2b*) embryos at 10, 15 and 20 hpf to overactivate FGF or BMP signaling throughout somitogenesis and concomitantly treated these embryos with either the BMP inhibitor DMH1 (12 µM) or the FGF inhibitor PD166866 (8 µM), respectively, between 10.5 and 28 hpf. Analysis of *nkx2.4b* expression in 28 hpf embryos showed that FGF overactivation could not rescue the loss of thyroid specification caused by DMH1 treatment (Fig. S5B data not shown). Conversely, the thyroid suppressive effect of the prolonged FGF inihbiton was partially rescued by repeated overactivation of BMP signaling (Fig. S5B). Collectively, these data are consistent with a model in which FGF signaling has permissive whereas BMP signaling has superordinate inductive capacity to initiate *nkx2.4b* expression in foregut endoderm.

## Discussion

Current models of thyroid development hold that a concurrent induction of *PAX8* and *NKX2-1*, or their teleost functional paralogs *pax2a* and *nkx2.4b*, is the initiating event of thyroid specification (Nilsson and Fagman, 2017; Porazzi et al., 2009; Szinnai, 2014) (De Felice and Di Lauro, 2011). In this study, we generated novel insights into the developmental dynamics of zebrafish thyroid cell specification permitting us to formulate a refined model of early thyroid cell differentiation.

First, we observed that the thyroid transcription factors Pax2a and Nkx2.4b are sequentially induced in the anterior foregut endoderm. Specifically, we identify a novel population of Pax2a+ endodermal cells emerging several hours before the onset of Nkx2.4b expression in the thyroid anlage region. We describe these early Pax2a+ cells as endodermal thyroid progenitors to distinguish them from subsequently evolving Pax2a+/Nkx2.4b+ cells which we refer to as lineage-committed thyroid precursors in accordance with previously proposed terminology (De Felice and Di Lauro, 2011).

The existence of an equivalent endodermal progenitor cell population expressing Pax8 prior to the onset of Nkx2-1 had not been previously reported in any of the commonly used model species. However, in ongoing preliminary studies we recently could confirm that a similar, supernumerary pool of Pax8+ endodermal cells exists during murine embryonic foregut development before the onset of Nkx2-1 expression (Opitz, R., unpublished observations). Although more studies in other species are needed, these findings lead us to predict that the molecular mechanisms of zebrafish thyroid specification reported here are likely conserved among vertebrates.

Our data indicate that thyroid lineage commitment is a multistep process where patterning processes initially yield a spatially restricted domain of Pax2a+ cells within the anterior foregut endoderm and a subsequent differentiation event results in the emergence of Pax2a+/Nkx2.4b+ thyroid precursors. This model is supported not only by the earlier appearance of Pax2a relative to Nkx2.4b expression but also by our observation that Nkx2.4b induction occurs exclusively in Pax2a+ endodermal cells. Another key finding of our studies was that the number of Pax2a+ cells populating the prospective thyroid field exceeds the final number of thyroid precursors contained in the thyroid bud. Throughout all stages of thyroid anlage formation (from 23/24 to 28 hpf), a variable number of Pax2a+ cells did not co-express Nkx2.4b and our temporal analyses suggest that these Pax2a+/Nkx2.4b-cells will not contribute to the thyroid primordium at later stages. We therefore conclude that Pax2a expression alone does not determine thyroid cell fate in zebrafish, but that thyroid lineage commitment requires co-expression of Pax2a and Nkx2.4b.

Over time, the initially supernumerary Pax2a+ cell population diminishes in size, and by thyroid bud stage, strong Pax2a expression was only maintained in cells co-expressing Nkx2.4b. Given that endodermal cells with gradually decreasing Pax2a+ immunolabeling remained well integrated in the ventral foregut epithelium, an explanation could be that, in the absence of Nkx2.4b expression, these initially Pax2a+ cells assume alternative cell fates within the pharyngeal epithelium after down-regulating Pax2a expression. Apoptosis cannot be ruled out, even though we have no supporting data.

According to this concept, multiple mechanisms seem to regulate endodermal Pax2a expression during early thyroid development. Early stimulatory signals, likely related to the general foregut endoderm patterning, induce Pax2a expression in a small medio-lateral stripe of the pharyngeal endoderm but these promoting effects appear transient. Additional regulatory mechanisms, linked to the co-expression of Nkx2.4b are required to stabilize Pax2a expression specifically in the thyroid cell lineage. Such a model of Pax2a regulation is in accordance with phenotypic studies of Nkx2.1/Nkx2.4b-deficient mouse ans zebrafish embryos showing that early *Pax8*/*pax2a* expression in the thyroid region is unaffected in Nkx2.1/Nkx2.4b-deficient embryos but *Pax8*/*pax2a* expression is subsequently lost in the absence of *Nkx2.1*/*nkx2.4b* function (Elsalini et al., 2003)(Parlato et al., 2004).

Although our current study showed that Nkx2.4b expression occurs exclusively in Pax2a+ endodermal cells, Pax2a expression itself is dispensable for Nkx2.4b induction in the zebrafish thyroid anlage (Trubiroha et al., 2018), as is Pax8 expression for Nkx2-1 induction in the murine thyroid anlage (Parlato et al., 2004). Thus, while Pax2a expression marks foregut endodermal cells that are competent to induce Nkx2.4b expression, absence of Pax2a expression does not impair this competence. Pax2a is, however, strictly required to maintain the thyroid fate and to promote further thyroid differentiation (Porreca et al., 2012; Trubiroha et al., 2018; Wendl et al., 2002).

The identification of distinct cell states during thyroid specification prompted us to examine the role of critical extrinsic factors such as FGF and BMP signaling in the spatiotemporal regulation of these processes. Prior to the appearance of Pax2a-expressing cells at 18 hpf, the prospective thyroid endoderm lacks detectable FGF and BMP signaling activity, an observation that aligns well with reported posteriorizing activities of FGF and BMP on endoderm patterning (Tiso et al., 2002) (Green et al., 2011). According to our refined model, induction of FGF signaling within the foregut endoderm is critical for the early onset of Pax2a expression. This contention is supported by several observations. First, Pax2a+ cells of the foregut endoderm are characterized by enhanced intracellular FGF signaling whereas adjacent Pax2a-negative endoderm lacks such FGF signaling activity. Second, blockade of FGF signaling diminishes the number of Pax2a+ cells in the thyroid field whereas late somitogenesis overactivation of FGF signaling causes induction of Pax2a expression in an increased number of endodermal cells. Finally, the proposed direct FGF regulation of early Pax2a expression in zebrafish foregut endoderm mirrors the capacity of FGF to induce high levels of PAX8 expression in human anterior foregut endoderm (Dye et al., 2015).

However, globally enhanced FGF signaling did not cause ectopic patches of Pax2a expression outside the thyroid field region in zebrafish embryos indicating that FGF alone has not sufficient instructive activity to convert neighboring endoderm into progenitors of the thyroid cell lineage. This conclusion is in accord with a lack of ectopic thyroid specification after grafting of FGF-soaked beads in the foregut region of zebrafish embryos (Wendl et al., 2007). It remains currently unclear if the spatial restriction of FGF effects on Pax2a expression is due to a cellular competence set by early endoderm patterning cues or if FGF signaling acts permissively together with other signaling cues (Haerlingen et al., 2019; Kurmann et al., 2015; Porazzi et al., 2012; Porreca et al., 2012; Rurale et al., 2020; Wendl et al., 2007).

One such additional pathway is BMP signaling (Haerlingen et al., 2019; Kurmann et al., 2015; Serra et al., 2017). In our model of sequential thyroid fate acquisition, the exclusive BMP pathway reporter expression in *nkx2.4b*-expressing cells clearly positions BMP activity as a key signaling cue regulating *nkx2.4b* expression in Pax2a+ endodermal progenitors. Our experimental data also indicate that BMP signaling has roles beyond the timely regulation of endodermal *nkx2.4b* induction because (1) BMP signaling can augment the FGF-dependent Pax2a induction, (2) ectopic expansion of the thyroid anlage is only seen upon concurrent overactivation of both BMP and FGF signaling at late somitogenesis stages and (3) inhibition of BMP signaling results in reduced endodermal Pax2a expression.

Our BMP signaling reporter experiments strongly suggest that promoting actions of BMP activity on Pax2a expression might be relayed via effects in adjacent non-endodermal tissue such as pre-cardiac and cardiac mesoderm. Previous studies have demonstrated that BMP signaling is active in the anterior LPM near the thyroid-forming endoderm and that BMP signaling is required for normal differentiation of these mesodermal derivatives (de Pater et al., 2012; Prummel et al., 2019). One appealing hypothesis, that needs confirmation, is that BMP activity is required for the proper development of the adjacent mesoderm, which in turn is responsible for the timely generation of signals (i.e. FGFs) that act on the endoderm to initiate Pax2a expression.

BMP-mediated patterning of mesoderm development is also likely involved in the unique anatomical constellation resulting from short-term overactivation of BMP signaling at the onset of segmentation. Here, the dramatic caudal expansion of the thyroid anlage was closely associated with the presence of caudally elongated strands of cardiac mesoderm bilaterally flanking a dysmorphic endoderm. Under the assumption that cardiac mesoderm-derived FGF and BMP signals act as critical signaling cues for thyroid specification at late somitogenesis, the latter phenotype illustrates the roles of BMP in mesoderm patterning and endodermal thyroid induction.

By integrating the various experimental findings, our revised model postulates that the midline positioning of the thyroid anlage stems from the intersection of two regional specification events. Spatial coordinates for thyroid specification along the AP axis would be determined by anterior LPM-derived patterning cues (including FGF ligands) resulting in a foregut endoderm region with a medial-to-lateral distribution of Pax2a+ cells at a defined AP level. This field of Pax2a+ cells has a competence for BMP-induced Nkx2.4b expression but only Pax2a+ cells closest to the cardiac mesoderm forming the apical pole of the heart tube might receive sufficiently strong BMP stimulation for Nkx2.4b induction. Thus, the midline positioning of the apical pole of the heart tube restricts thyroid precursor specification to more medial positions within the broader Pax2a+ field. Such a dual step specification process provides a very informative concept explaining the various thyroid phenotypes arising in response to cardiac maldevelopment (Haerlingen et al., 2019; Vandernoot et al., 2020).

## Acknowledgments

The authors would like to thank Jean-Marie Vanderwinden and staff from the Light Microscopy Facility for confocal microscopy assistance.

## Author Contributions

B.H. performed most of the experiments under the supervision of R.O.. B.H. and R.O. wrote the manuscript. A.M., R.O., I.V. and, P.G. helped to perform the BMP and FGF activity inhibition and mapping experiments. B.H., R.O., and S.C. designed the experiments and interpreted the data with the help of A.T.. M.S., P.G., A.T. and S.C. edited the manuscript.

## Declaration of Interests

The authors declare no conflicts of interest.

## Competing Interest Statement

The authors have declared no competing interest.

## STAR Methods

### LEAD CONTACT AND MATERIALS AVAILABILITY

Further information and requests for resources and reagents should be directed to and will be fulfilled by the Lead Contact, Sabine Costagliola. This study did not generate new unique reagents, plasmids or animal lines.

### EXPERIMENTAL MODEL AND SUBJECT DETAILS

Zebrafish husbandry and experiments with all transgenic lines will be performed under standard conditions as per the Federation of European Laboratory Animal Science Associations (FELASA) guideline, and in accordance with institutional (Université Libre de Bruxelles (ULB)) and national ethical and animal welfare guidelines and regulation, which were approved by the ethical committee for animal welfare (CEBEA) from the Université Libre de Bruxelles (protocols 578N-579N).

Embryos were collected within 30 minutes after natural spawning, raised in embryo medium at 28.5°C and staged according to hours post fertilization (hpf) as described (Haerlingen et al., 2019; Kimmel et al., 1995). Media were supplemented with 0.003% 1-phenyl-2-thiourea (PTU) from 24 hpf onwards to prevent pigmentation. Embryos were enzymatically dechorionated by incubation in embryo medium containing 0.6 mg/mL pronase at room temperature(Haerlingen et al., 2019). For live analyses and before sampling or fixation, embryos were anesthetized in medium containing 0.02% tricaine. Embryos were fixed in 4% paraformaldehyde (PFA) solution in phosphate-buffered saline (PBS) overnight at 4°C with gentle agitation.

The following zebrafish lines were used: pigmentless *casper* strain (White et al., 2008), Et(*nkx2.4b*:YFP) (CLGY576) (Ellingsen et al., 2005), Tg(*hsp70*:ca-fgfr1,*cryaa*:DsRed)^pd1^ abbreviated as Tg(*hsp70*:ca-fgfr1) (Marques et al., 2008), Tg(*hsp70*:*bmp2b*)^fr13^ (Chocron et al., 2007), Tg(−5.0*sox17*:EGFP)^ha01^ abbreviated as Tg(*sox17:*EGFP) (Mizoguchi et al., 2008), Tg(*myl7*:EGFP)^twu26^ (Huang et al., 2003), Tg(*dusp6*:d2EGFP)^pt6^ (Molina et al., 2007), Tg(BRE-AAVmlp:dmKO2)^mw40^ abbreviated as Tg(BRE:dmKO2) (Collery and Link, 2011).

## METHOD DETAILS

### Small molecule treatments

Stock solutions of 20 mM PD166866 and 10 mM DMH1 were prepared in DMSO and aliquots were stored at −20°C until use. Treatment solutions were freshly prepared by diluting stock solutions in embryo medium and kept at 28.5°C. The final test concentrations of 8 µM PD166866 and 12 µM DMH1 were selected for strong inhibitory activity based on results of our previous to have optimal inhibition effects with virtually no toxic effects (Haerlingen et al., 2019). Embryos treated with 0.1% DMSO served as a vehicle control group for all small molecule treatment experiments. All solutions were supplemented with 0,01 mg/mL methylene blue to prevent bacterial growth and, if treatment runs until 24 hpf or later and if embryos are intended for whole mount *in situ* hybridization (WISH), with 0,003% 1-phenyl-2-thiourea (PTU).

Two hours before treatment initiation, embryos were staged and stage-matched embryos were pooled and randomly allocated to the different experimental treatments into 60mm Petri dishes, 40-50 maximum per Petri dish. Treatment periods are indicated in the text and the corresponding figures. All treatments were performed in duplicates. All incubations of embryos in small molecule solutions were performed at 28.5°C in the dark. After completion of small molecule treatment, embryos were washed three times with fresh embryo medium before transfer into clean 96 mm Petri dishes filled with fresh embryo medium.

### Heat shock treatment

Embryos for heat shock (HS) experiments were obtained by crossing heterozygotic founder fish of the Tg(*hsp70l*:ca-fgfr1) or Tg(*hsp70l*:*bmp2b*) lines with homozygotic founder fish of the indicated reporter lines Et(*nkx2.4b*:YFP), Tg(*myl7*:EGFP) and Tg(*sox17*:EGPF). Stage-matched embryos obtained from such crosses were pooled and randomly allocated to the differentially timed HS experimental groups. HS was typically applied at either 10 hpf (early somitogenesis), 15 hpf (mid somitogenesis) or 20 hpf (late somitogenesis). Some experiments involved repeated HS treatment of embryos at 10, 15 and 20 hpf. HS experiments were performed at least in duplicates. For HS treatment, embryo media were entirely removed and replaced by pre-warmed embryo medium (40°C) and embryos were incubated at 40°C for 30 min. After the HS period, warm media were entirely removed and replaced by fresh embryo medium and embryos were further raised under standard conditions at 28.5°C. In some experiments, HS treatments were combined with small molecule inhibitor treatments. Here, pre-warmed embryo medium containing the indicated small molecule inhibitors was used for the 40°C incubation to ensure continuous inhibitor exposure.

Heat-shocked embryos were genotyped by PCR after completion of whole mount *in situ* hybridization as previously described (Shin et al., 2007). Embryos carrying the *hsp70l*:*bmp2b* transgene were identified by amplifying a segment overlapping on the *hsp70l* promoter and the *bmp2b* coding sequence (forward primer: 5’-CATGTGGACTGCCTATGTTCATC-3’; reverse primer: 5’-GAGAGCGCGGACCACGGCGAC-3’). Embryos carrying the *hsp70l*:*ca-fgfr1* transgene were identified by amplifying a portion of the DsRed coding sequence that is part of the transgenic cassette (forward primer: 5’-CTCCAAGGCCTACGTGAAGCAC-3’; reverse primer: 5’-CACGGGCTTCTTGGCCTTG-3’). Overactivation of BMP signaling in embryos obtained from outcrosses of heterozygotic Tg(*hsp70l:bmp2b*) founder fish caused a characteristic tail phenotype in about 50% of heat-shocked embryos when HS treatment was performed at 10 hpf. Genotyping showed that the tail phenotype clearly segregated with the presence of the *hsp70l:bmp2b* allele in all embryo clutches analyzed. If indicated, this phenotypic trait was used in some experiments to distinguish WT and *hsp70l:bmp2b* allele carriers when genotyping was difficult to perform.

### Single and dual-color whole-mount in situ hybridization

DNA templates for synthesis of *nkx2.4b* (previously *nkx2.1a*, ZDB-GENE-000830-1), *tg* (ZDB-GENE-030519-1), *hhex* (ZDB-GENE-980526-299), *pax2a* (ZDB-GENE-990415-8) and *foxa2* (ZDB-GENE-980526-404) riboprobes were generated by PCR essentially as described (Opitz et al., 2011). Riboprobes labeled with digoxigenin (DIG), fluorescein (FLU) or dinitrophenol (DNP) were prepared as described (Opitz et al., 2011) using DIG RNA labeling kit (Roche), FLU RNA labeling kit (Roche) or DNP-UTP (Perkin Elmer).

The whole-mount *in situ* hybridization (WISH) protocol was based on Thisse and Thisse (C. Thisse and B. Thisse, 2008) with modifications described in (Haerlingen et al., 2019; Opitz et al., 2011). Briefly, PFA-fixed embryos were washed three times for 10 min with PBST followed by gradual dehydration through a series of methanol/PBST solutions (25, 50, 75, 100%) and stored in 100% methanol at −20°C for at least 24 hours. Following gradual rehydration in PBST, embryos were permeabilized by short-term proteinase K treatment, post-fixed for 20 min in 4% PFA and rinsed in PBST. Hybridization of embryos was performed at 65°C essentially as described (Opitz et al., 2011).

Single color staining of embryos hybridized with DIG-labelled riboprobes was performed using anti-DIG antibody conjugated to alkaline phosphatase (Roche, 1:6000) and BM purple (Roche) as alkaline phosphatase substrate as described (Opitz et al., 2011). Dual color stainings were performed as described (Haerlingen et al., 2019). Briefly, following hybridization of embryos with multiple differentially labeled riboprobes, we first detected the DIG-labelled *nkx2.4b* riboprobe with anti-DIG antibody (1:6000) and BM purple staining in 28 hpf embryos and the DNP-labelled *tg* riboprobe with anti-DNP antibody (Vector Laboratories, 1:500) and NBT/BCIP (Roche) staining in 55 hpf embryos. Stained specimens were washed in PBST, shortly cleared in 100% methanol to remove unspecific background staining, rehydrated in PBST, and incubated in HCl-glycine solution (pH 2.2) for two times 5 min to effectively remove antibodies used for the first round of staining. Embryos were then incubated in anti-FLU antibody conjugated to alkaline phosphatase (Roche, 1:2000) and expression of *myl7* in 28 and 55 hpf embryos was then revealed by Fast Red (Sigma) staining. Embryos were shortly post-fixed in 4% PFA, rinsed in PBST and transferred to 95% glycerol. Whole mount imaging of WISH-stained specimen was performed using a Leica DFC420C camera mounted on a Leica MZ16F stereomicroscope.

### Whole-mount immunofluorescence

Whole-mount immunofluorescence (WIF) was essentially performed as previously described (Opitz et al., 2011). Briefly, PFA-fixed embryos were washed several times with PBS containing 0.1% Tween20 (PBST). Embryos were mildly permeabilized by proteinase K treatment, washed with PBST and incubated for 4-6 hours in blocking buffer (PBST containing 10 mg/mL bovine serum albumin, 4% horse serum, 1% DMSO and 0,8% Triton X-100) at room temperature with gentle agitation. Embryos were then incubated in blocking buffer containing primary antibodies overnight at 4°C. After removal of primary antibody solutions, embryos were extensively washed in PBST and incubated in blocking buffer containing secondary antibodies overnight at 4°C. After removal of secondary antibody solutions, embryos were washed in PBST and incubated for three days in PBST containing Hoechst nuclear dye (1:7500 dilution) to counterstain cell nuclei. After several PBST washes, embryos were shortly post-fixed in 4% PFA for 20 min at room temperature and stored in PBST at 4°C until whole mount imaging or vibratome sectioning.

The following antibodies were used: rabbit anti-Pax2a (1:250); mouse anti-E Cadherin (1:200); chicken anti-GFP (1:1000, also recognizes d2EGFP and YFP); rabbit anti-monomeric destabilized Kusabira Orange 2 (dmKO2, 1:250); mouse anti-dmKO2 (1:200); Cy3-conjugated donkey anti-rabbit IgG (1:250); Cy3-conjugated donkey anti-mouse IgG (1:250); Alexa Fluor 647-conjugated donkey anti-rabbit IgG (1:250); Alexa Fluor 647-conjugated donkey anti-mouse IgG (1:250); Alexa Fluor 488-conjugated goat anti-chicken IgG (1:250); Alexa Fluor 488-conjugated donkey anti-chicken IgG (1:250); Alexa Fluor 647-conjugated donkey anti-goat IgG (1:250) (see Key Resources Table for suppliers and references).

For whole mount imaging, WIF-stained specimen were embedded in 1% low melting agarose (Lonza) on Fluoro-Dish glass bottom dishes (World Precision Instruments). Images were acquired using a Leica DFC7000T camera mounted on a Leica M165FC stereomicroscope and LAS X software (Leica).

### Confocal image acquisition & cell counting

Confocal microscopic analyses of the thyroid region of zebrafish embryos was performed based on vibratome sections of WIF-stained specimen. After WIF staining, embryos were embedded in 7% low melting agarose and serial 100 µm thick sections were cut on a Leica VT1000S vibratome. Sections were mounted in Glycergel (Dako) on custom-made #1.5 72×26 mm glass slides. Confocal images were acquired using an LSM510 confocal microscope (Zeiss) and Zen 2010 D software (Zeiss). For quantitative analyses of cell numbers, confocal *Z*-stacks covering the whole thyroid region were acquired and the number of cells expressing specific markers was determined manually by analyzing consecutive *Z*-stacks. Images were processed on ImageJ/Fiji with a 1-pixel radius Minimum filter to remove artefacts.

## STATISTICAL ANALYSIS

For statistical analyses of cell number measurements, pairwise comparisons were conducted using parametric *t*-test with a Welch’s correction and a two-tailed *P* value (95%). Statistical analyses were performed using Graph-Pad Prism7 (GraphPad, San Diego, CA). Differences were considered significant at *P* < 0.05.

---

**Fig. S1:**
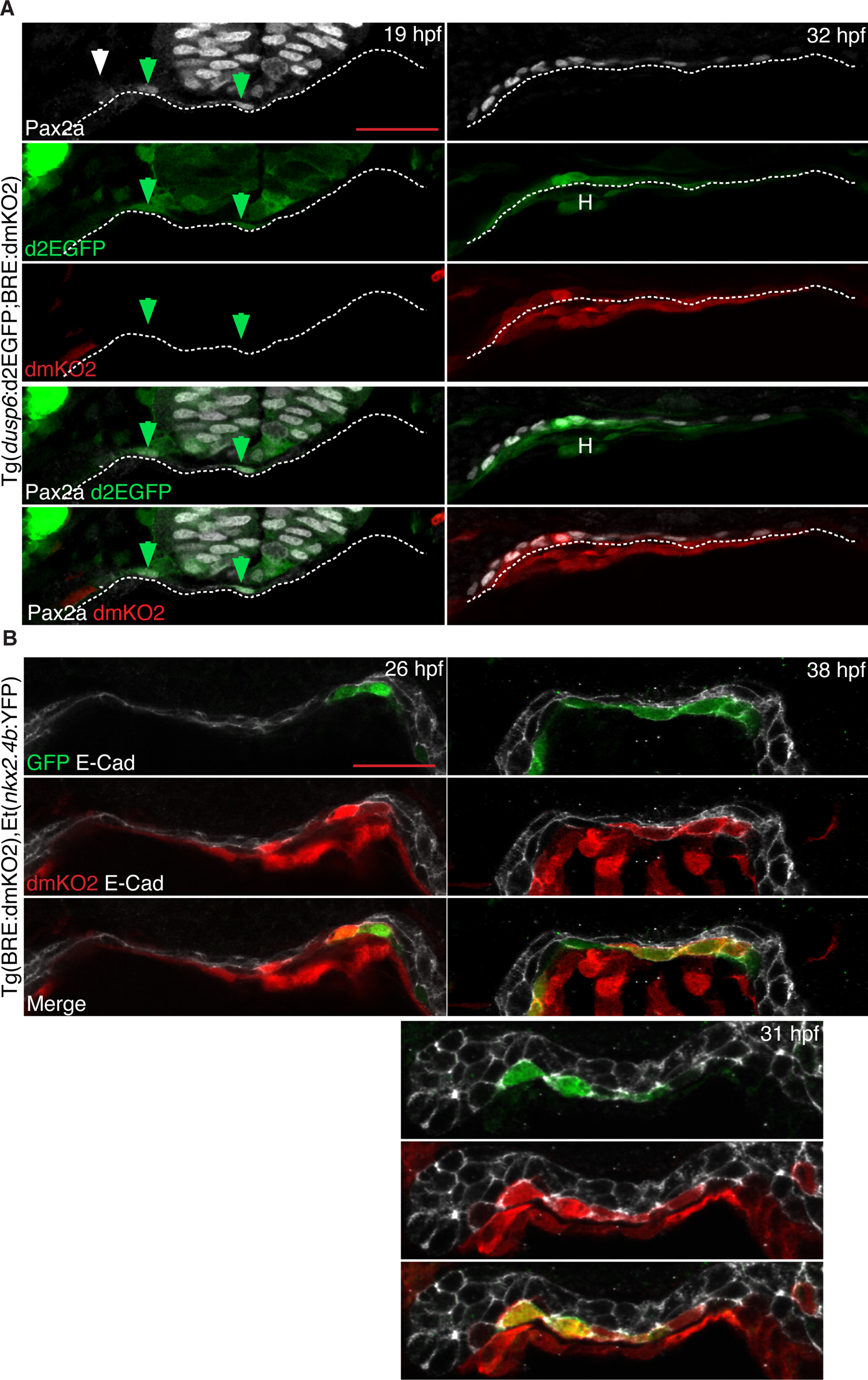
FGF and BMP activity in thyroid precursors and progenitors. (A) Immunofluorescence of Pax2a, d2GFP and dmKO2 expression in thyroid region of Tg(*dusp6*:d2EGFP;BRE:dmKO2) double transgenic embryos. Confocal images of frontal sections are shown. Dashed line depicts border between endodermal cell layer and ventral foregut mesenchyme. Note the absence of BMP signaling reporter expression (dmKO2) in endoderm and Pax2a+ cells at 19 hpf and the spatially restricted dmKO2 expression in Pax2a+ endodermal cells adjacent to cardiac mesoderm at 32 hpf. Arrowheads in left panel highlight Pax2a+ cells expressing the FGF signaling reporter d2EGFP. H: heart. Scale bar: 25 µm. (B) Immunofluorescence of E-Cadherin (ECad), dmKO2 and YFP expression in thyroid region of Tg(BRE:dmKO2);Et(*nkx2.4b:*YFP) double transgenic embryos. Confocal images of frontal sections are shown. E-cadherin labels the foregut endoderm. Note that YFP expression is restricted to a few endodermal cells expressing the BMP signaling reporter dmKO2. Scale bar: 25 µm.

**Fig. S2:**
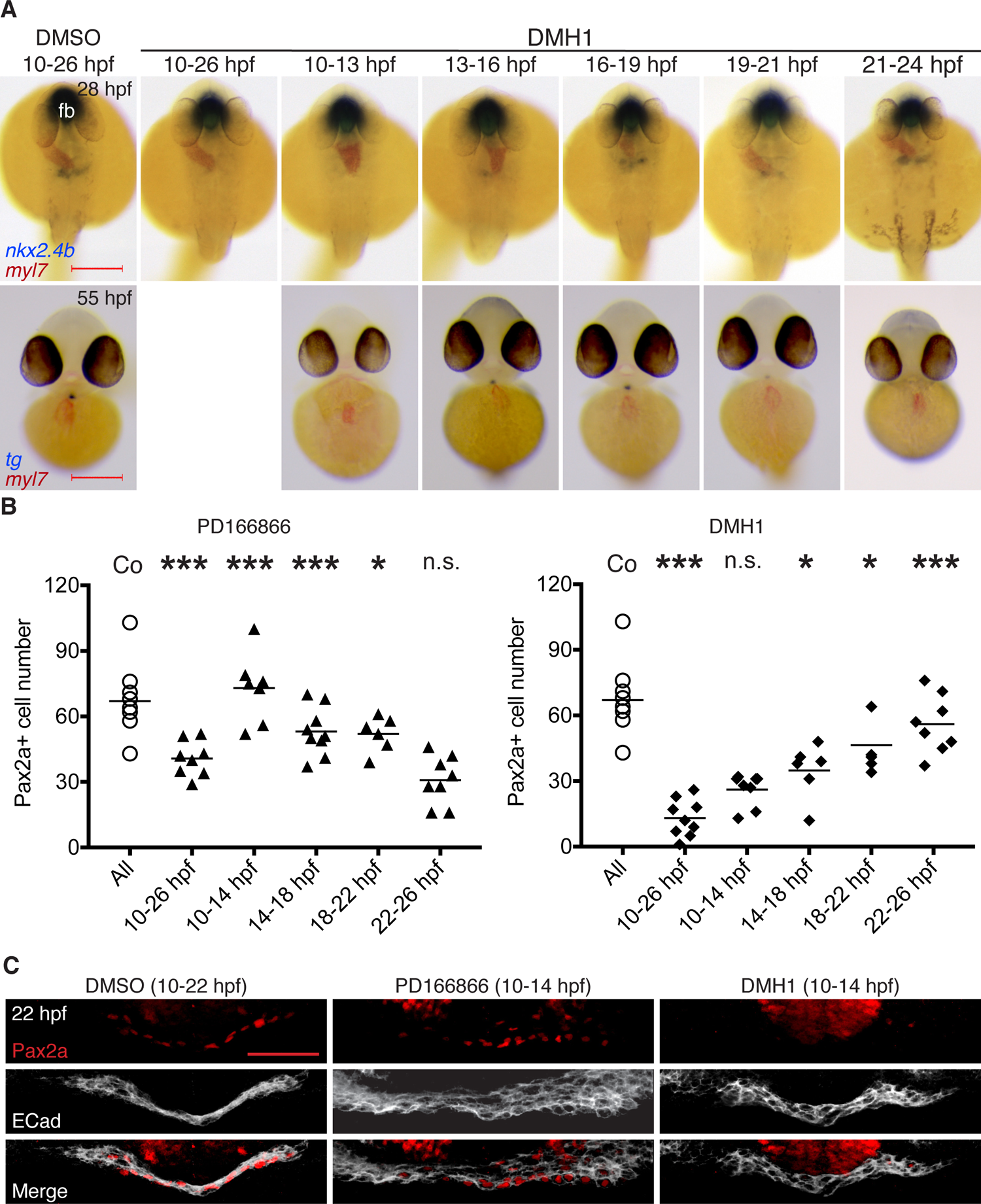
Impaired thyroid specification after short BMP and FGF inhibitions. (A) Major thyroid phenotypes recovered following DMH1 treatment during the indicated periods. Thyroid development was assessed by *in situ* hybridization of *nkx2.4b* at 28 hpf (upper panel, dorsal views, anterior to the top) and *tg* at 55 hpf (lower panel, ventral views, anterior to the top). Co-staining with *myl7* riboprobe shows localization of cardiac tissue. Note the severe repression of thyroid marker expression due to DMH1 treatment at early somitogenesis (10-13 hpf) and during thyroid specification (19-21 and 21-24 hpf) fb: forebrain. Scale bar: 200 µm. (B) Quantification of the number of foregut endoderm cells expressing Pax2a in 28 hpf embryos following short-term treatment with PD166866 (8 µM) or DMH1 (12 µM) for the indicated periods. Embryos treated with 0.1% DMSO served as a vehicle control group. Values obtained in individual embryos are shown and bars depict mean values for each experimental group. Asterisks denote significant differences between individual treatments and the control (n.s. no significant difference, * *P*<0.05, *** *P*<0.001). Note that values of the control group represent pooled control data from the various treatment periods. (C) Three-dimensional reconstruction of Pax2a expression domain in the thyroid region of 22 hpf embryos treated during early somitogenesis with inhibitors of FGF signalling (8 µM PD166866) or BMP signaling (12 µM DMH1). Confocal images were acquired from frontal vibratome sections. E Cadherin (ECad) labels the foregut endoderm. Note the loss of Pax2a expression in response to DMH1 treatment between 10 and 14 hpf. Scale bar: 25 µm.

**Fig. S3:**
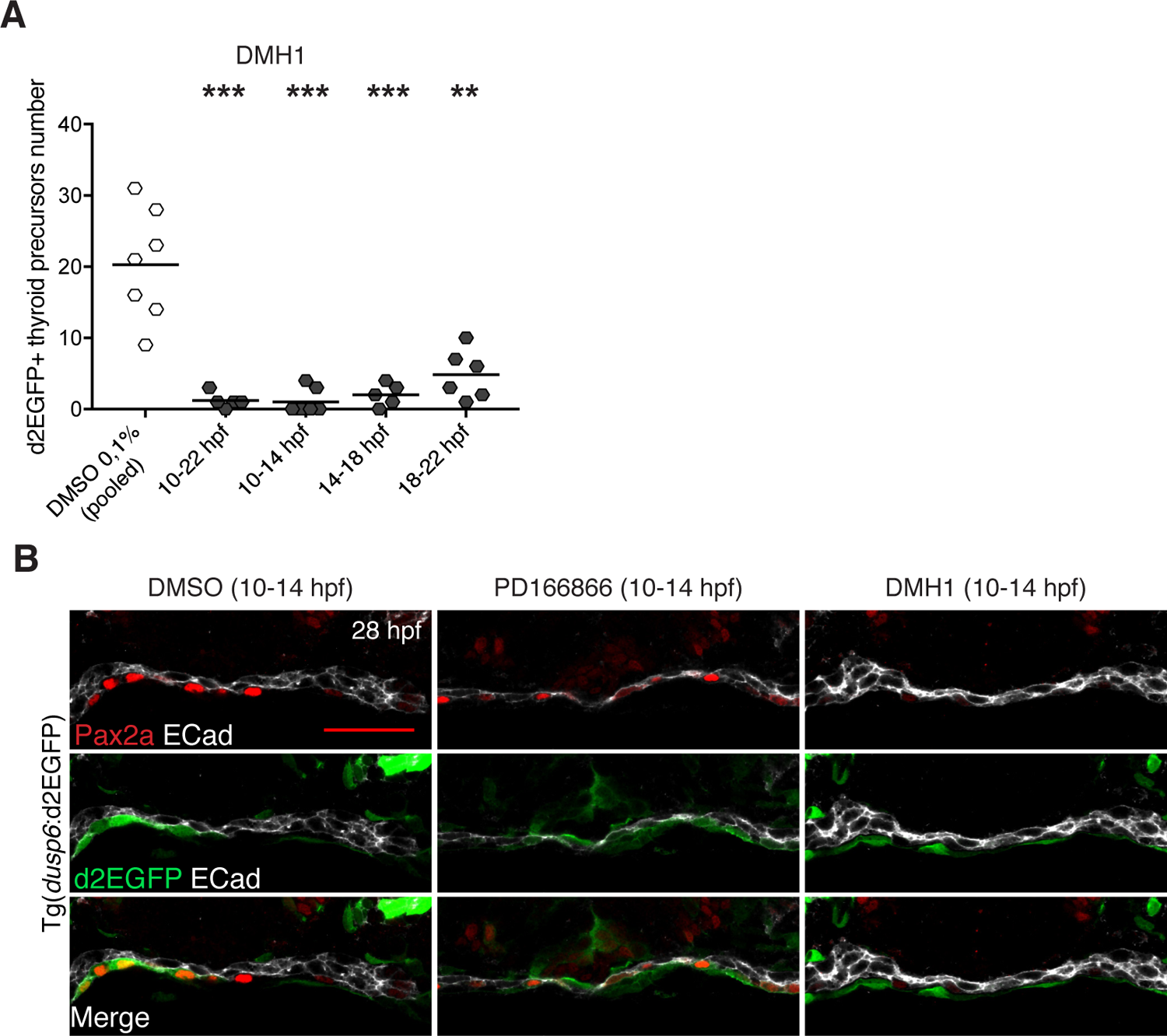
Early somitogenesis BMP inhibition affects FGF activity at thyroid specification stages. (A) Quantification of the number of foregut endoderm cells co-expressing Pax2a and the FGF signaling reporter d2EGFP following short-term inhibition of BMP signaling. Embryos were treated with either 12 µM DMH1 or 0.1% DMSO (vehicle control) for the indicated periods. Values obtained in individual 28 hpf embryos are shown and bars depict mean values for each experimental group. Asterisks denote significant differences between individual treatments and the control (** *P*<0.01, *** *P*<0.001). Note that values of the control group represent pooled control data from the various treatment periods. (B) Immunofluorescence of Pax2a, E-cadherin (ECad) and d2EGFP expression in thyroid region of 28 hpf Tg(*dusp6*:d2EGFP) transgenic embryos following short-term inhibition of FGF or BMP signaling at early somitogenesis. Embryos were treated from 10-14 hpf with 8 µM PD166866 or 12 µM DMH1 to block FGF and BMP signaling, respectively. Confocal images of frontal sections are shown. Note the absence of FGF signaling reporter expression in foregut endoderm following BMP inhibitor treatment. Scale bar: 25 µm.

**Fig. S4:**
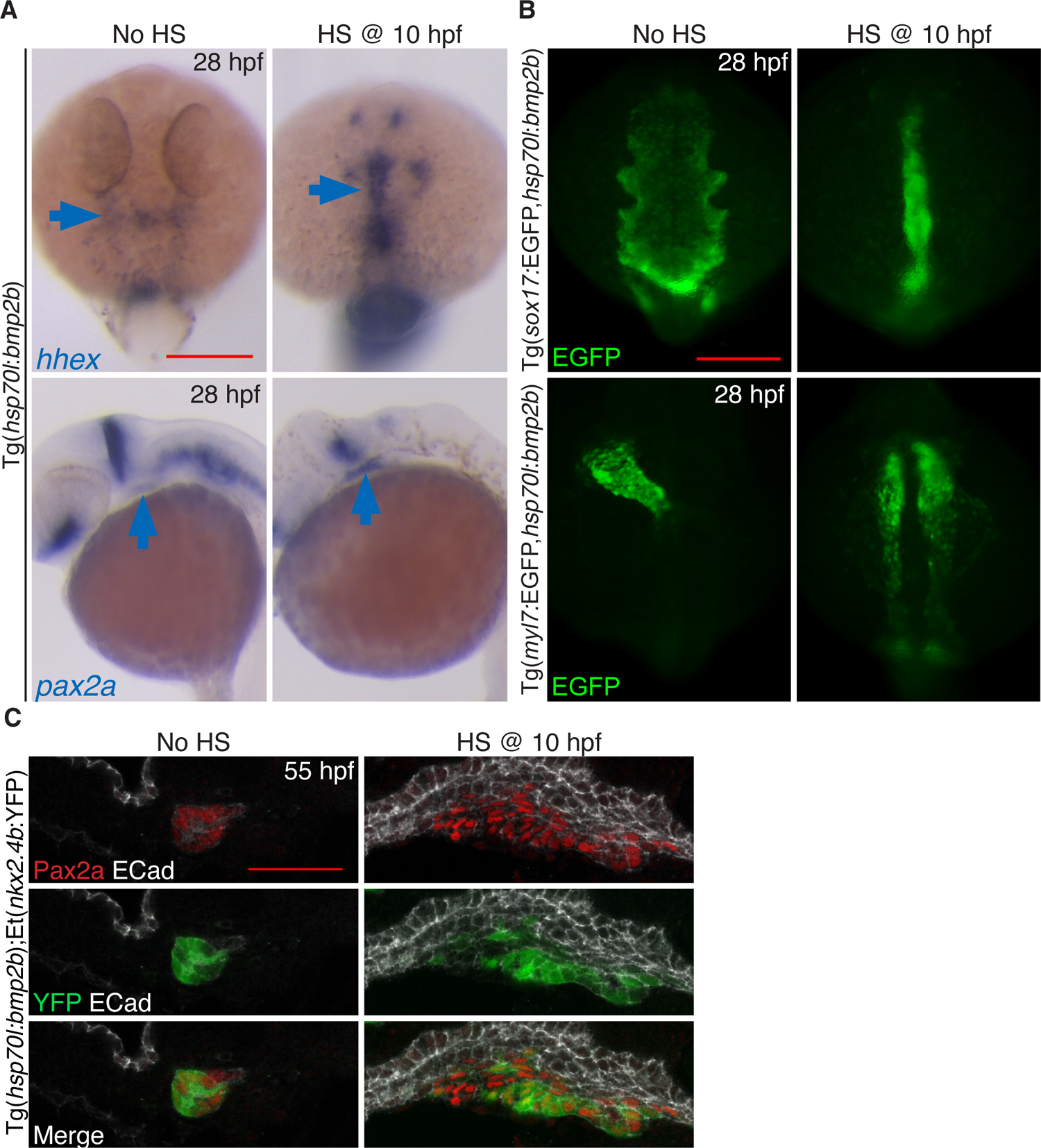
Global over-activation of BMP signaling during early somitogenesis promotes massive expansion of thyroid precursors. (A) Whole mount *in situ* hybridization of 28 hpf embryos for early thyroid markers *hhex* (dorsal views, anterior to the top) and *pax2a* (lateral views, anterior to the left) showed an aberrant enlargement of the thyroid anlage when heat shock (HS) treatment of Tg(*hsp70l*:*bmp2b*) embryos was applied at early somitogenesis. Note the caudally expanded expression thyroid domains of *hhex* and *pax2a*. Scale bar: 200 µm. (B) Whole mount immunofluorescence of GFP reporter expression in 28 hpf Tg(*hsp70l*:*bmp2b,sox17*:EGFP) and Tg(*hsp70l*:*bmp2b,myl7*:EGFP) embryos revealed severe maldevelopment of foregut endoderm (upper panel) and cardiac mesoderm (lower panel) if HS treatment was applied at 10 hpf. Note the abnormal rod-like appearance of the foregut endoderm and the failure of midline fusion and heart tube assembly of cardiomyocytes. Dorsal views are shown, anterior to the top. Scale bar: 200 µm. (C) Immunofluorescence of Pax2a, E-Cadherin (ECad) and YFP expression in thyroid region of 55 hpf double transgenic Tg(*hsp70l*:*bmp2b*),Et(*nkx2.4b*:YFP) embryos demonstrates a massive enlargement of the thyroid primordium when overactivation of BMP signalling was induced by HS treatment at 10 hpf. Confocal images of sagittal sections are shown, anterior is to the left. Note the large amount of Pax2a+/YFP+ thyroid tissue ventral to the pharyngeal epithelium in heat-shocked embryos and the contrasting small compact thyroid primordium present in 55 hpf controls. Scale bar: 25 µm.

**Fig. S5:**
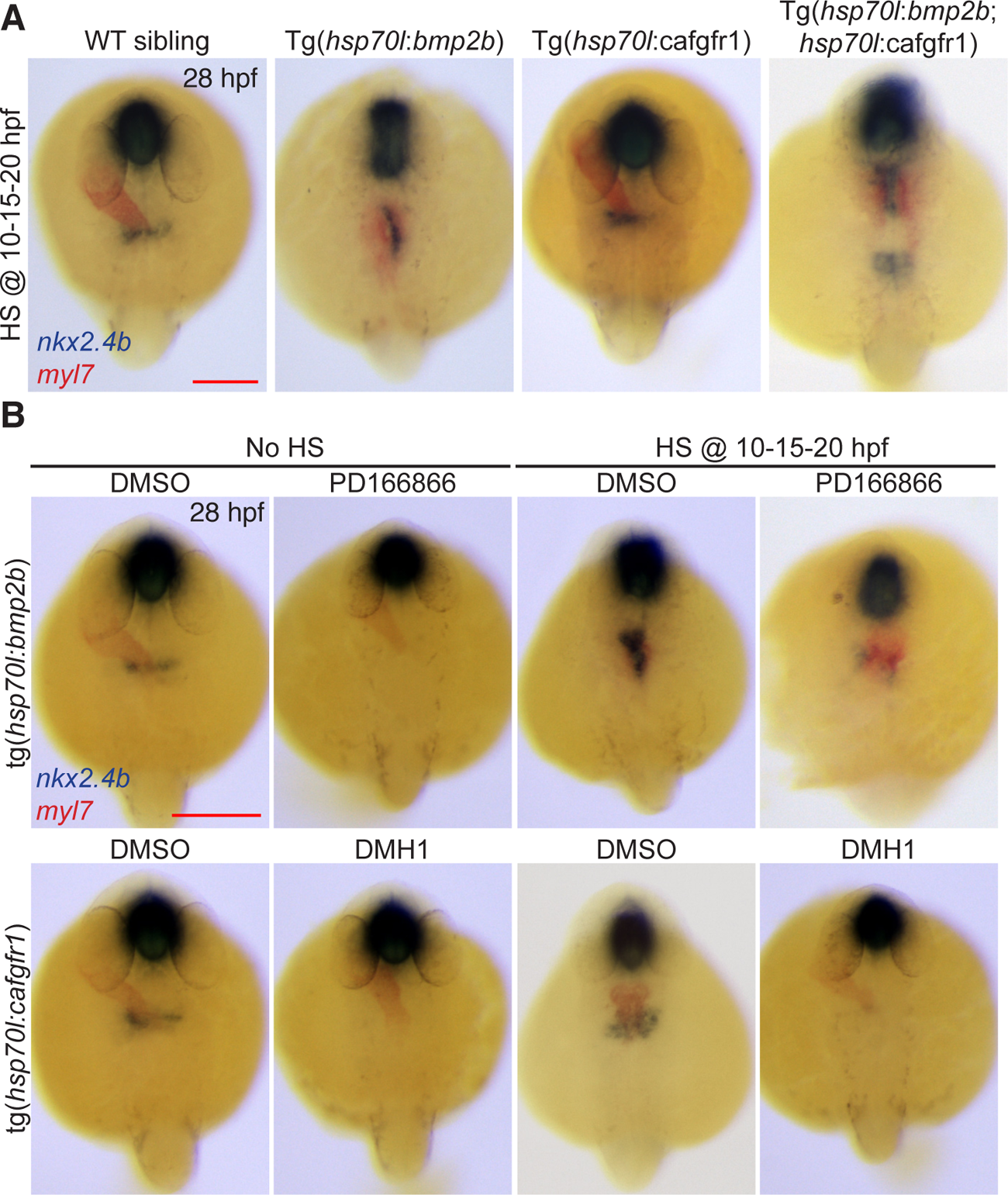
Combined BMP and FGF activation causes amplified thyroid enlargement. (A) Major thyroid phenotypes recovered following repeated heat shock-induced overactivation of BMP, FGF or FGF and BMP signaling throughout somitogenesis. Thyroid development was assessed by dual-color *in situ* hybridization of *nkx2.4b* and *myl7*. Note the presence of abnormal, caudally expanding *nkx2.4b* expression domain following overactivation of BMP signaling alone as well as following concurrent overactivation of both FGF and BMP signaling. Combined overactivation of FGF and BMP signaling induced additional ectopic *nkx2.4b* expression domains at more caudal positions. Dorsal views, anterior to the top. fb: forebrain. Scale bar: 200 µm. Upper panel shows that repeated HS-induced overactivation of BMP signaling can partially rescue the thyroid agenesis phenotype caused by PD166866 inhibition of FGF signaling. Embryos were treated with either 8 µM PD166866 or 0.1% DMSO (vehicle control) from 10 to 24 hpf. HS of Tg(*hsp70l*:*bmp2b*) embryos was repeatedly applied at 10, 15 and 20 hpf. Bottom panel shows that repeated HS-induced overactivation of FGF signaling cannot rescue the thyroid agenesis phenotype caused by DMH1 inhibition of BMP signaling. Embryos were treated with either 12 µM DMH1 or 0.1% DMSO (vehicle control) from 10 to 24 hpf. HS of Tg(*hsp70l*:*cafgfr1*) embryos was repeatedly applied at 10, 15 and 20 hpf. Scale bar: 200 µm.

